# RAF1 scaffold integrity shapes chemogenetic degradation outcomes in KRAS-driven lung cancer

**DOI:** 10.64898/2026.06.23.733960

**Authors:** Laura de-la-Puente-Ovejero, Ana Domostegui, Inés María García-Pérez, Gonzalo Aizpurua, Lucía Lomba-Riego, Pilar Ximenez-Embún, Cristina Mayor-Ruiz, Mariano Barbacid, Sara García-Alonso

## Abstract

Scaffold integrity is essential for the activity of proteins that function through protein-protein interactions rather than catalytic output. RAF1 exemplifies this duality: although it is a bona fide kinase and a core component of the MAPK cascade, its tumor-promoting role is largely kinase-independent, relying instead on scaffold-mediated suppression of apoptosis. Genetic *Raf1* ablation in KRAS-driven lung adenocarcinoma mouse models induces tumor regression without systemic toxicity, making it an attractive candidate for targeted protein degradation. Chemogenetic systems like the dTAG platform are widely used for preclinical target validation. Here, we generated a dTAG-RAF1 mouse model and showed that pharmacological degradation is efficient and systemically well tolerated, but fails to reproduce the tumor regression observed upon genetic *Raf1* ablation. Mechanistically, the N-terminal FKBP12^F36V^ tag (dTAG) perturbs the RAF1 interactome, including scaffold associations with apoptotic regulators, thereby blunting the phenotypic consequences of its degradation. These results establish scaffold integrity as a determinant of chemogenetic system fidelity and argue that degradation tools must be validated at the functional level, not only for target elimination, before assessing their therapeutic relevance.

## Introduction

Mutations in the RAS gene family occur in approximately 30% of all human cancers, with KRAS being the most frequently altered isoform (Prior *et al*, 2012). These mutations are particularly prevalent in pancreatic, colorectal, and lung adenocarcinomas (LUAD), and are strongly associated with poor prognosis (Kodaz, 2017). Despite the central role of KRAS in tumorigenesis, it was long considered “undruggable” (Moore & Malek, 2021). The discovery of a small, druggable pocket in the KRAS^G12C^ isoform (Ostrem *et al*, 2013) enabled the development of selective inhibitors such as sotorasib and adagrasib, both of which have received FDA approval (Skoulidis *et al*, 2021; Jänne *et al*, 2022; Yaeger *et al*, 2023). However, clinical responses remain limited by modest efficacy in some settings and by the emergence of adaptive and acquired resistance mechanisms (Arbour *et al*, 2023; Strickler *et al*, 2023). As a result, research has expanded toward pan-RAS inhibitors, combination therapies, and targeting of downstream signaling nodes, particularly the RAF/MEK/ERK cascade, a critical axis in RAS-driven malignancies (Barrambana *et al*, 2025).

Among RAF kinases, RAF1 (also known as CRAF) has emerged as a relevant downstream effector and potential therapeutic target in KRAS-driven LUAD. Conditional deletion of *Raf1* in genetically engineered mouse models (GEMMs) induces tumor regression without major toxicity in normal tissues, indicating that RAF1 is essential for tumor maintenance but dispensable for adult tissue homeostasis (Sanclemente *et al*, 2018). Moreover, *Raf1* ablation alone is sufficient to promote tumor regression in this setting, with co-targeting of related kinases or receptor tyrosine kinases providing no additional benefit (de-la-Puente-Ovejero *et al*, 2026). This contrasts with MEK or ERK inhibition, which, although effective in preclinical tumor models, results in systemic toxicity due to the essential roles of these kinases in normal physiology (Drosten & Barbacid, 2020). Interestingly, the tumor-suppressive effects of RAF1 do not arise from the loss of canonical MAPK signaling, but rather from the activation of pro-apoptotic programs normally repressed by RAF1 kinase-independent functions (Sanclemente *et al*, 2021). Here, we refer to these non-catalytic, protein-interaction-mediated activities as RAF1 scaffold functions. In this context, scaffold integrity refers to the ability of RAF1 to preserve the interaction surfaces and protein associations required for its non-catalytic functions.

These features make RAF1 a challenging target for conventional pharmacological approaches. Unlike BRAF, which has a relatively accessible ATP-binding pocket suitable for type I/II kinase inhibitors (Cook & Cook, 2021), the RAF1 kinase domain is conformationally shielded by the HSP90-CDC37 chaperone complex (García-Alonso *et al*, 2022), limiting accessibility to typical small molecules. Furthermore, pharmacological inhibition of RAF1 catalytic activity is unlikely to fully reproduce the biological consequences of RAF1 loss, given that its tumor-promoting effects in KRAS-driven LUAD are largely non-catalytic. Together, these observations support approaches aimed at eliminating RAF1 protein rather than inhibiting its kinase activity.

Targeted protein degradation (TPD) is a pharmacological strategy that enables the modulation of protein abundance. “Degraders” such as proteolysis targeting chimeras (PROTACs), molecular glues, and other bifunctional molecules exploit the ubiquitin-proteasome system to induce selective elimination of proteins of interest (POIs) (Békés *et al*, 2022). By recruiting an E3 ubiquitin ligase to a target protein, degraders promote ubiquitination and subsequent proteasomal degradation of the POI. Unlike occupancy-driven inhibitors, TPD has the potential to suppress both catalytic and non-catalytic protein functions by removing the target protein itself (Paiva & Crews, 2019). This makes degradation-based approaches particularly attractive for proteins like RAF1, whose oncogenic functions extend beyond enzymatic activity.

Nevertheless, applying TPD to scaffold proteins presents specific challenges. First, degrader molecules must engage structurally accessible surfaces on the POI, which can be difficult when the target is embedded in large regulatory complexes, such as RAF1 within the RAF1-HSP90-CDC37 (RHC) complex (García-Alonso *et al*, 2022). Second, the biological outcome of degradation depends not only on efficient target elimination, but also on whether loss of the protein reproduces the relevant functional consequences of genetic ablation. Third, some proteins, such as RAF1, currently lack high-affinity selective ligands; in those cases, chemogenetic degradation systems are often used for initial therapeutic validation (Mayor-Ruiz & Winter, 2019). Widely used systems such as the degradation TAG (dTAG) system (Nabet *et al*, 2018) enable the rapid and selective elimination of otherwise undruggable proteins. However, the required N-or C-terminal fusion of chemogenetic tags to the POI can sometimes disturb native conformation, subcellular localization, or interaction networks, thereby complicating biological interpretation (Taylor *et al*, 2025).

Indeed, emerging evidence suggests that proteins carrying chemogenetic tags may exhibit altered function even prior to degradation. Several studies have shown that chemical degradation of these chimeric proteins may yield weaker or divergent phenotypes compared with genetic ablation (Barsoum *et al*, 2023; Yenerall *et al*, 2023). Therefore, it is essential to validate degradation tools not only at the level of target elimination, but also at the level of functional equivalence to the endogenous protein.

In this study, we used dTAG-RAF1 as a model to evaluate both the feasibility of inducible protein degradation *in vivo* and the functional fidelity of chemogenetic tagging applied to an oncogenic protein. We generated a conditional KRAS-driven LUAD mouse model in which endogenous *Raf1* was replaced by a chemically degradable FKBP12^F36V^-tagged (dTAG) variant. Using this system, we assessed systemic tolerability, tumor response, MAPK signaling, apoptosis, and RAF1-associated protein interactions following degradation with the VHL-recruiting PROTAC dTAG^V^-1m. Unexpectedly, although dTAG-RAF1 was efficiently degraded and systemic RAF1 elimination was well tolerated, the tagged protein was not fully functionally equivalent to endogenous RAF1. The N-terminal FKBP12^F36V^ tag selectively impaired apoptosis-associated interactions while preserving MAPK-associated signaling, thereby limiting the ability of the model to reproduce the effects of genetic *Raf1* ablation. As an exploratory resource, we also annotated reported RAF1 ubiquitination sites onto the RHC complex to identify solvent-exposed lysines within the native chaperone-bound state. Collectively, our findings emphasize that chemogenetic degradation systems for scaffold proteins require validation beyond degradation efficiency, particularly when the relevant biological activity depends on protein-protein interactions rather than catalytic output.

## Results

### N-terminal FKBP12^F36V^-RAF1 preserves MAPK signaling but alters KRAS-driven tumor progression

To establish a chemogenetic system suitable for inducible RAF1 degradation, we first evaluated whether fusion of dTAG to RAF1 interfered with RAF1 function. N-and C-terminal tagged RAF1 constructs were tested in RAF-deficient (RAFless: *Araf*^L/Y^*; Braf*^L/L^*; Raf1*^L/L^) cells lacking endogenous ARAF, BRAF, and RAF1 expression following 4-hydroxytamoxifen (4-OHT) treatment. Whereas the N-terminally dTAG-RAF1 fusion partially restored colony formation, the C-terminal tag insertion failed to rescue proliferation (Fig EV1), indicating that C-terminal tagging compromises RAF1 functionality. Based on these observations, the N-terminal configuration was selected for subsequent studies.

Using homologous recombination, we generated a knock-in mouse model in which the endogenous *Raf1* allele was replaced by a chemically degradable FKBP12^F36V^-tagged version (dTAG-RAF1), enabling pharmacological control of RAF1 abundance (Fig 1A). Expression analysis confirmed that dTAG-RAF1 was produced at levels comparable to endogenous RAF1 at both the protein and mRNA levels (Fig 1B-C).

**Fig 1.**
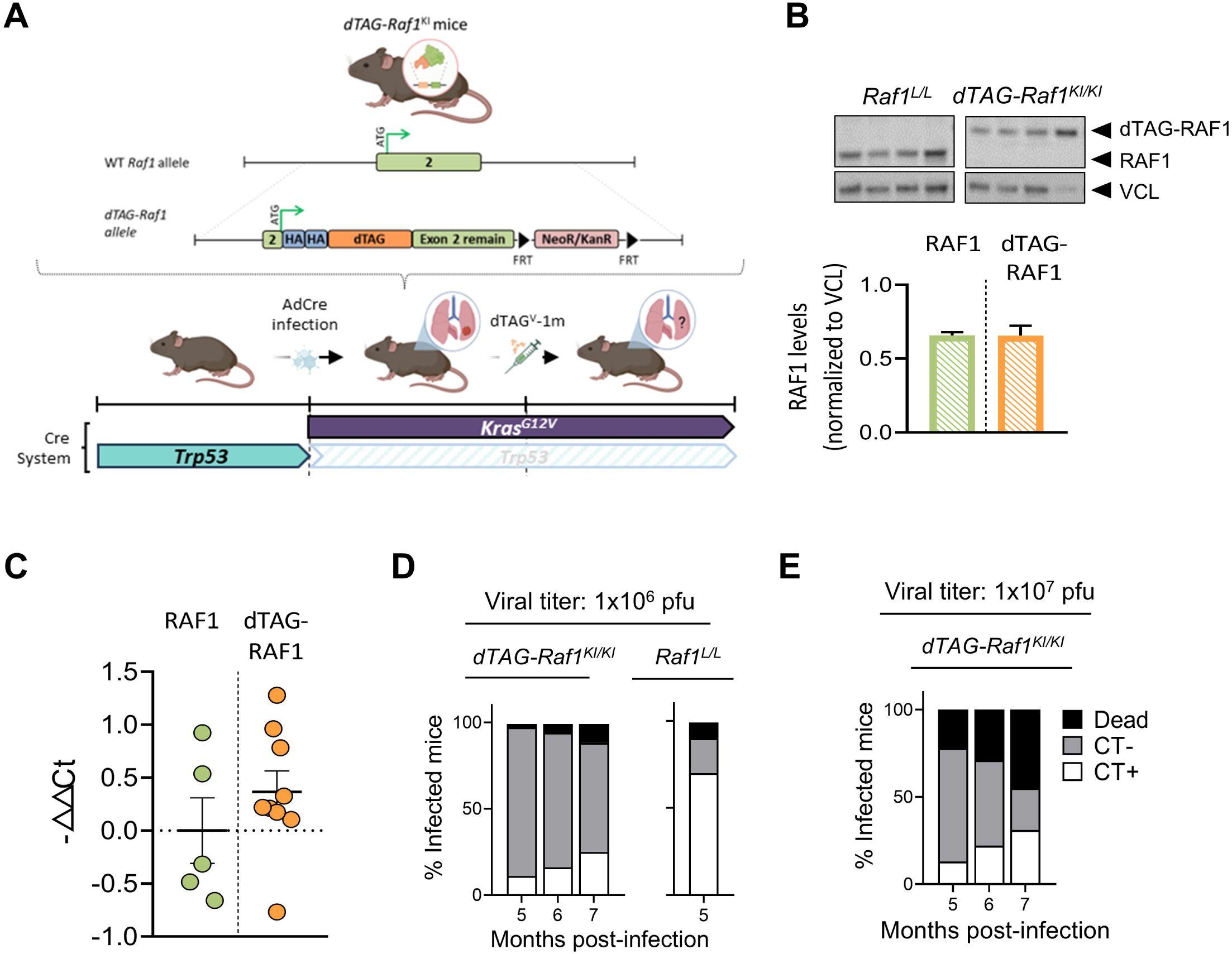
Expression of dTAG-RAF1 delays KRAS-driven lung tumor development. (A) Schematic of the *dTAG-Raf1*^KI^ mouse model and experimental design. Exon 2 of *Raf1* (green) was modified to include a double HA tag (blue) followed by the FKBP12^F36V^ sequence (orange), with a NeoR/KanR selection cassette (pink) flanked by FRT sites inserted downstream. *dTAG-Raf1*^KI/KI^ mice were infected intranasally with AdCre to induce *Kras*^G12V^ expression and *Trp53* deletion in lung epithelial cells, generating LUAD. Tumor-bearing mice were subsequently treated with dTAG^V^-1m to achieve pharmacological RAF1 elimination at the protein level. Solid-colored boxes indicate expressed alleles; striped boxes represent ablated genes. (B) Western blot analysis and quantification of RAF1 protein levels in *Raf1*^L/L^ and dTAG*-Raf1*^KI/KI^ mice, normalized to VCL. (C) qPCR quantification of *Raf1* mRNA from the same genotypes. Data are presented as −ΔΔCt relative to the housekeeping control. Data in B and C are presented as mean ± SEM. Statistical comparisons were performed using unpaired Student’s t-test; no significant differences were detected. (D-E) Tumor onset in *Kras*^+/G12V^; *Trp53*^-/-^; dTAG*-Raf1*^KI/KI^ and control (*Kras*^+/G12V^; *Trp53*^-/-^; *Raf1*^L/L^) mice following infection with 1×10^6^ or 1×10_ pfu AdCre. Tumor detection by CT scan is shown as a percentage of total infected mice, classified as tumor-free (CT-negative, grey), tumor-bearing (CT-positive, white), or deceased (black) at each time point. Panel D includes 87 dTAG*-Raf1*^KI/KI^ mice and 55 *Raf1*^L/L^ mice from a historical cohort, shown at 5 months post-infection for reference; panel E includes 45 dTAG-*Raf1*^KI/KI^ mice.

To determine whether expression of the tagged protein affected KRAS-driven tumorigenesis, *Kras*^+/LSLG12V^; *Trp53*^L/L^; *dTAG-Raf1*^KI/KI^ mice were infected intranasally with AdCre and monitored by computerized tomography (CT) imaging. Unexpectedly, tumor onset was significantly delayed relative to mice expressing endogenous WT RAF1. While *Kras*^+/LSLG12V^; *Trp53*^L/L^; *Raf1*^L/L^ mice reproducibly develop detectable lung tumors within approximately 5 months after infection, only a minority of *Kras*^+/LSLG12V^; *Trp53*^L/L^; *dTAG-Raf1*^KI/KI^ mice showed CT-detectable lesions even after prolonged follow-up (Fig 1D). Increasing the viral titer tenfold did not enhance tumor penetrance and instead increased toxicity without improving tumor burden (Fig 1E).

These observations indicated that expression of dTAG-RAF1 partially alters RAF1-dependent tumor-promoting functions *in vivo* despite preserved expression levels.

As an exploratory resource for future RAF1 degrader development, we also annotated reported RAF1 ubiquitination sites onto available RAF1 structural models. Direct experimental detection of RAF1 ubiquitinated peptides after HSP90 destabilization and proteasome inhibition was technically challenging, likely due to poor recovery from insoluble material (Fig EV2A-B). Integration of large-scale ubiquitinome datasets with curated repositories and structural mapping identified K462 and K575 as the most solvent-exposed lysines among the structurally resolved RAF1 ubiquitination sites (Fig EV2C-E). These observations do not predict degrader-productive ubiquitination but provide structural annotation of accessible RAF1 surfaces within the native RHC complex.

### Systemic degradation of dTAG-RAF1 is well tolerated in adult mice

We next assessed whether pharmacological elimination of dTAG-RAF1 produced systemic toxicity *in vivo*. Adult tumor-free *Kras*^+/LSLG12V^; *Trp53*^L/L^; *dTAG-Raf1*^KI/KI^ mice (not infected with AdCre) were treated daily for 30 days with the VHL-recruiting degrader dTAG^V^-1m (Martín *et al*, 2026) (40 mg/kg, i.p.). In parallel, a second compound, IGP002 (a PROTAC analogue with improved solubility and metabolic stability) was also evaluated.

Western blot analysis confirmed efficient degradation of dTAG-RAF1 in lung and liver following dTAG^V^-1m treatment (Fig 2A). Although IGP002 (Martín *et al*, 2026) also induced dTAG-RAF1 degradation, it additionally led to off-target degradation of endogenous FKBP12 and was therefore excluded from further analysis.

**Fig 2.**
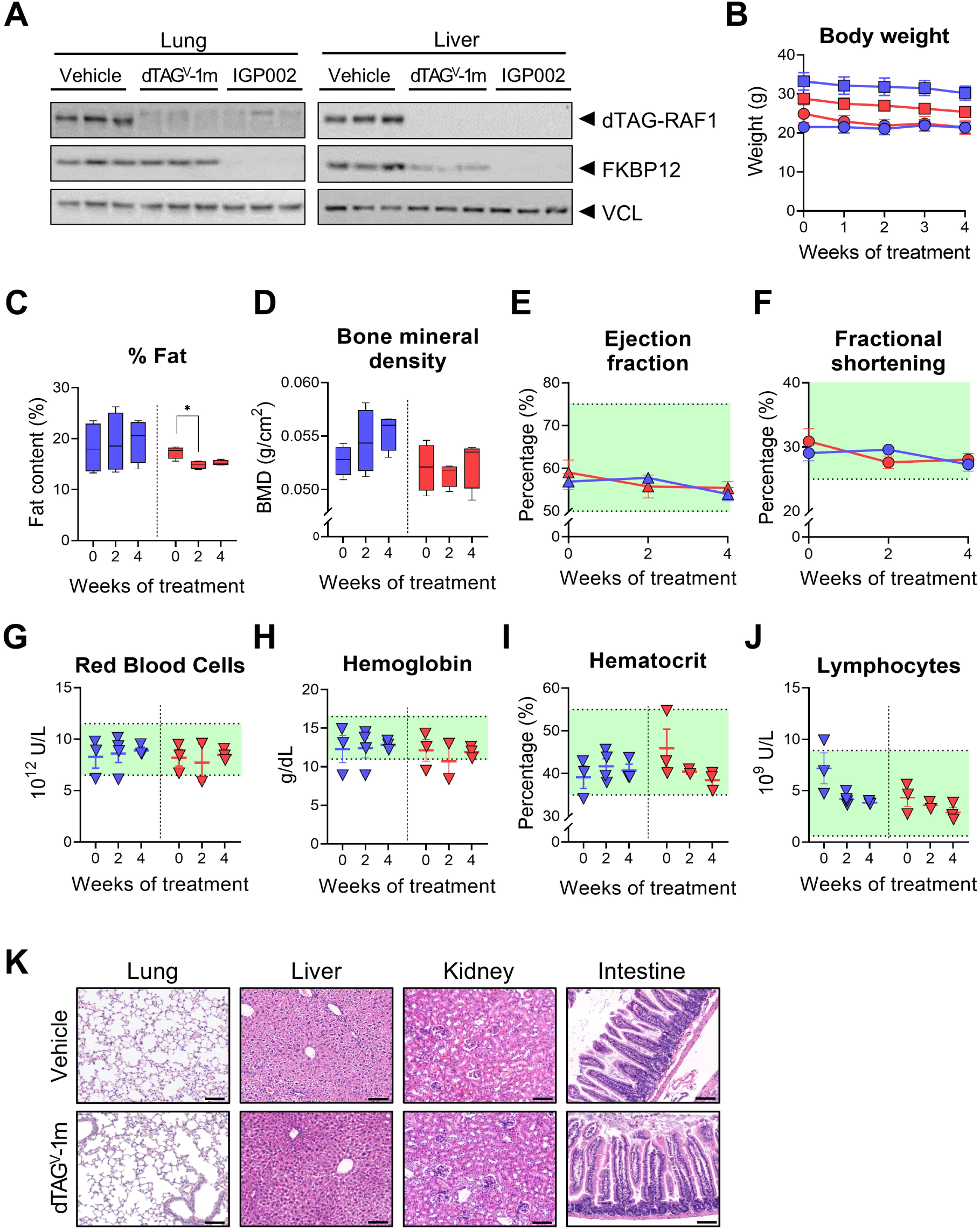
Systemic dTAG^V^-1m-induced degradation of RAF1 is well tolerated *in vivo*. (A) Western blot showing effective degradation of dTAG-RAF1 across major tissues following daily dTAG^V^-1m and IGP002 administration (40 mg/kg) for 30 days. (B) Body weight profiles of vehicle-and dTAG^V^-1m-treated *Kras*^+/LSLG12V^; *Trp53*^L/L^; *dTAG-Raf1*^KI/KI^ mice. Males and females are represented by squares and circles, respectively. (C–D) Densitometric analysis of body fat percentage and bone mineral density using DEXA scanning. (E–F) Echocardiographic assessment of ejection fraction and fractional shortening. (G–J) Hematologic parameters, including RBC count, HGB, HCT, and lymphocyte percentage. Vehicle-and dTAG^V^-1m-treated groups are shown in blue and red, respectively. Green shaded areas in panels E–J indicate physiological reference ranges. (K) Representative H&E staining of major organs from vehicle-and dTAG^V^-1m-treated animals. Scale bars: 50 µm. Statistical comparisons were performed using paired Student’s t-test (n = 4 mice per group). * p < 0.05.

Comprehensive physiological characterization revealed no systemic toxicity associated with sustained dTAG-RAF1 degradation. Body weight remained stable in both dTAG^V^-1m-and vehicle-treated groups (Fig 2B). DEXA scans showed no significant changes in body composition or bone mineral density (Fig 2C-D). Echocardiographic assessment showed preserved ejection fraction and fractional shortening across all groups and time points (Fig 2E-F), indicating maintained cardiovascular integrity. Similarly, hematological parameters remained within physiological ranges, with no significant changes in erythroid or lymphoid populations (Fig 2G-J).

Histopathological examination of major organs, including lung, liver, kidney, and intestine, revealed no detectable pathological alterations or tissue damage following treatment (Fig 2K). No clinical signs of distress or treatment-associated morbidity were observed during the study.

Together, these findings indicate that systemic pharmacological degradation of dTAG-RAF1 is well tolerated in adult mice and support previous genetic evidence suggesting that RAF1 is dispensable for normal tissue homeostasis.

### Efficient degradation of dTAG-RAF1 does not recapitulate the tumor regression observed after genetic *Raf1* ablation

To evaluate the consequences of acute RAF1 elimination in established tumors, *Kras*^+/G12V^; *Trp53*^-/-^; *dTAG-Raf1*^KI/KI^ mice bearing CT-confirmed lung tumors (≥ 5 mm³) were treated daily with dTAG^V^-1m (40 mg/kg i.p.). Tumor growth was monitored longitudinally by CT every 15 days.

Western blotting and immunohistochemistry confirmed robust degradation of dTAG-RAF1 protein in tumor tissues after treatment (Fig 3A-B). However, despite efficient and sustained protein elimination, tumors failed to regress. Longitudinal CT imaging instead revealed continued tumor growth throughout the treatment period (Fig 3C). Linear mixed-effects modeling confirmed a statistically significant positive slope in tumor volume over time in dTAG^V^-1m-treated mice, indicative of ongoing tumor progression (Fig 3D).

**Fig 3.**
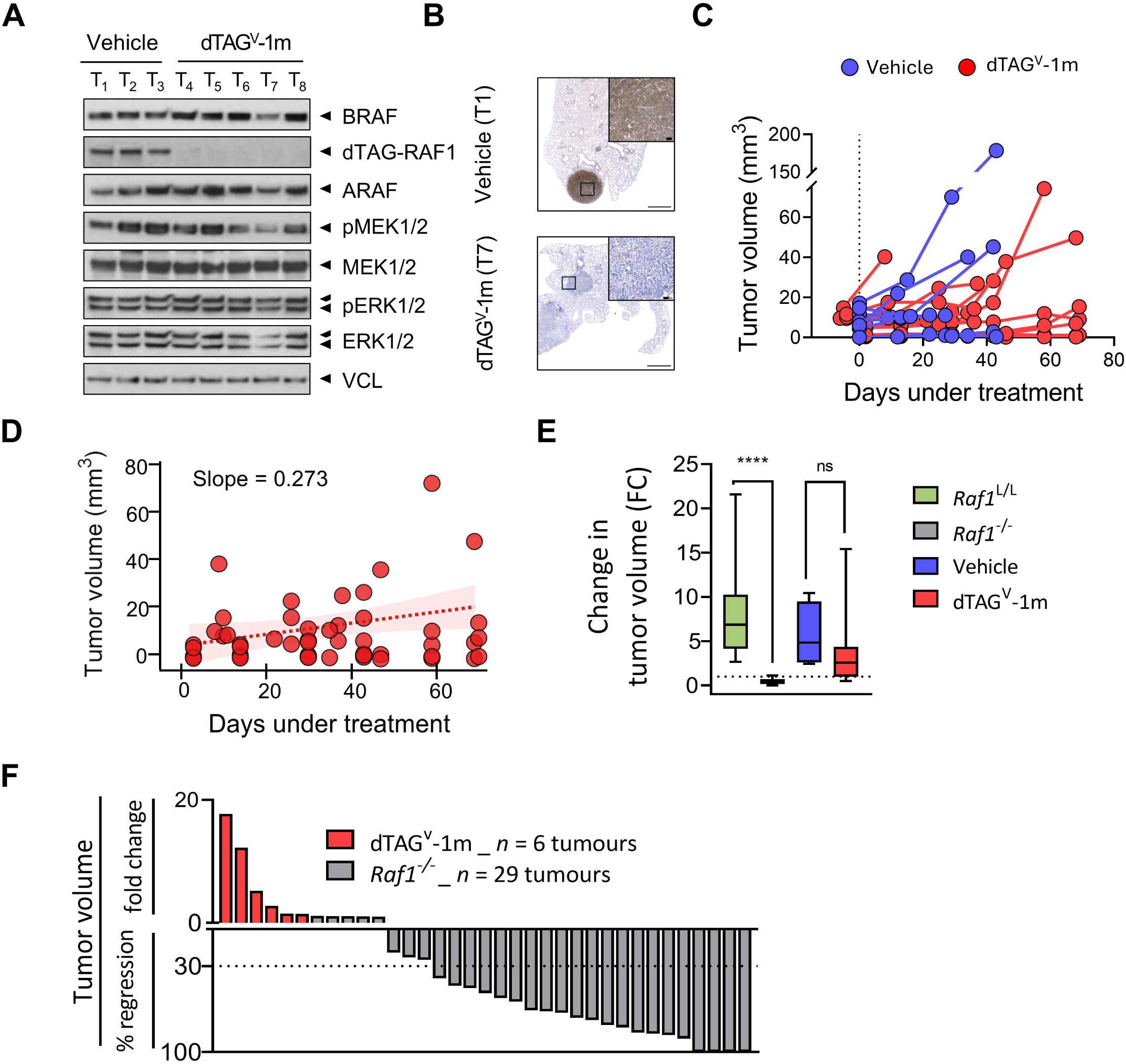
Efficient degradation of dTAG-RAF1 fails to induce tumor regression. (A) Western blot validation of dTAG-RAF1 degradation and downstream MAPK pathway in tumor samples. VCL was used as a loading control. (B) Representative RAF1 immunostaining of paraffin-embedded lung tumors from vehicle-and dTAG^V^-1m-treated mice. Scale bars: 1 mm (overview), 50 μm (zoom). (C) Longitudinal tumor volume measurements by CT imaging of individual tumors from vehicle-(blue; 6 mice, 7 tumors) and dTAG^V^-1m-treated (red; 7 mice, 13 tumors) groups. (D) Linear mixed-effects model of tumor growth over time. (E) Comparison of tumor responses to RAF1 elimination across models represented as FC in tumor volume, based on CT scans performed at the beginning and end of the study. Tumors from *Kras*^+/G12V^; *Trp53*^-/-^; *Raf1*^L/L^ (*n* = 44, green), *Kras*^+/G12V^; *Trp53*^-/-^; *Raf1*^-/-^ (*n* = 40, grey), vehicle-treated *Kras*^+/G12V^; *Trp53*^-/-^; *dTAG-Raf1*^KI/KI^ (*n* = 4, blue), and dTAG^V^-1m-treated *Kras*^+/G12V^; *Trp53*^-/-^; *dTAG-Raf1*^KI/KI^ (*n* = 9, red) mice are shown. Statistical significance was calculated according to the Mann-Whitney test. (F) Waterfall plot representing individual tumor volume changes in *Kras*^+/G12V^; *Trp53*^-/-^; dTAG*-Raf1*^KI/KI^ mice treated with dTAG^V^-1m (red) and *Kras*^+/G12V^; *Trp53*^-/-^; *Raf1*^-/-^ mice exposed to 4-OHT diet (grey) for 2 months. Comparison with historical *Kras*^+/G12V^; *Trp53*^-/-^; *Raf1*^L/L^ and *Kras*^+/G12V^; *Trp53*^-/-^; *Raf1*^-/-^ cohorts is presented for contextual reference; direct statistical comparison is limited by differences in experimental design, treatment agent, and cohort size. Statistical comparisons in E were performed using the Mann-Whitney test; tumor growth in D was assessed using a linear mixed-effects model. Significance thresholds: * p < 0.05, ** p < 0.01, *** p < 0.001, **** p < 0.0001.

To contextualize these findings, tumor responses were compared across *Kras*^+/G12V^; *Trp53*^-/-^; *Raf1*^L/L^, *Kras*^+/G12V^; *Trp53*^-/-^; *Raf1*^-/-^, vehicle-treated *Kras*^+/G12V^; *Trp53*^-/-^; *dTAG-Raf1*^KI/KI^, and dTAG^V^-1m-treated *Kras*^+/G12V^; *Trp53*^-/-^; *dTAG-Raf1*^KI/KI^ cohorts. The *Raf1*^L/L^ and *Raf1*^-/-^ data derived from historical cohorts generated in our laboratory. These data are consistent with the interpretation that pharmacological degradation of dTAG-RAF1 does not reproduce the tumor regression phenotype observed after genetic *Raf1* ablation (Fig 3E-F).

To complement these *in vivo* findings, primary LUAD cell lines derived from the same GEMM were established and treated *in vitro* with dTAG^V^-1m. Consistent with the *in vivo* response, dTAG-RAF1 degradation did not impair the proliferative capacity of these cells (Fig EV3A-B), in contrast to previous observations using WT RAF1-targeting shRNAs, which robustly inhibited colony formation (Sanclemente *et al*, 2018).

Collectively, these results demonstrate that efficient degradation of dTAG-RAF1 does not phenocopy the anti-tumor effects previously observed following genetic *Raf1* deletion.

### dTAG-RAF1 preserves MAPK signaling but selectively disrupts scaffold-associated interactions

To investigate why dTAG^V^-1m-mediated RAF1 degradation failed to induce tumor regression, we analyzed whether the dTAG protein retained the signaling and scaffold properties associated with endogenous RAF1.

In RAFless cells, both WT RAF1 and dTAG-RAF1 partially restored colony formation following elimination of endogenous RAF proteins (Fig EV3C). In addition, kinase assays revealed similarly low catalytic activity for WT and dTAG-RAF1 within the HSP90-CDC37 complex context (Fig EV3D), consistent with previous observations that RAF1 kinase activity is tightly regulated by chaperone association (Aizpurua *et al*, 2026).

Analysis of downstream MAPK signaling showed no major differences between WT and dTAG-RAF1 conditions. ERK phosphorylation remained detectable in tumors before and after dTAG^V^-1m treatment (Fig 3A), and confocal microscopy revealed comparable subcellular localization patterns for WT and dTAG-RAF1 (Fig EV3E).

Because RAF1-dependent tumor maintenance has been linked primarily to kinase-independent anti-apoptotic functions, we next examined apoptosis-associated responses. In contrast to the phenotype observed after genetic *Raf1* ablation, degradation of dTAG-RAF1 did not induce detectable accumulation of cleaved caspase-3 (cl.CASP3) (Fig 4A). In *Raf1*^L^ cells, treatment with etoposide or staurosporine produced only limited induction of apoptotic markers, as assessed by cleaved PARP (cl.PARP) and cl.CASP3. In contrast, cells expressing dTAG-RAF1 showed an increase in these markers following the same treatments (Fig 4B). Similar results were obtained when apoptosis was analyzed by flow cytometry (Fig 4C). Together, these data support a model in which *Raf1*^L^ retains anti-apoptotic functions, whereas the N-terminal tag in dTAG-RAF1 compromises scaffold integrity and sensitizes cells to apoptotic stimuli.

**Fig 4.**
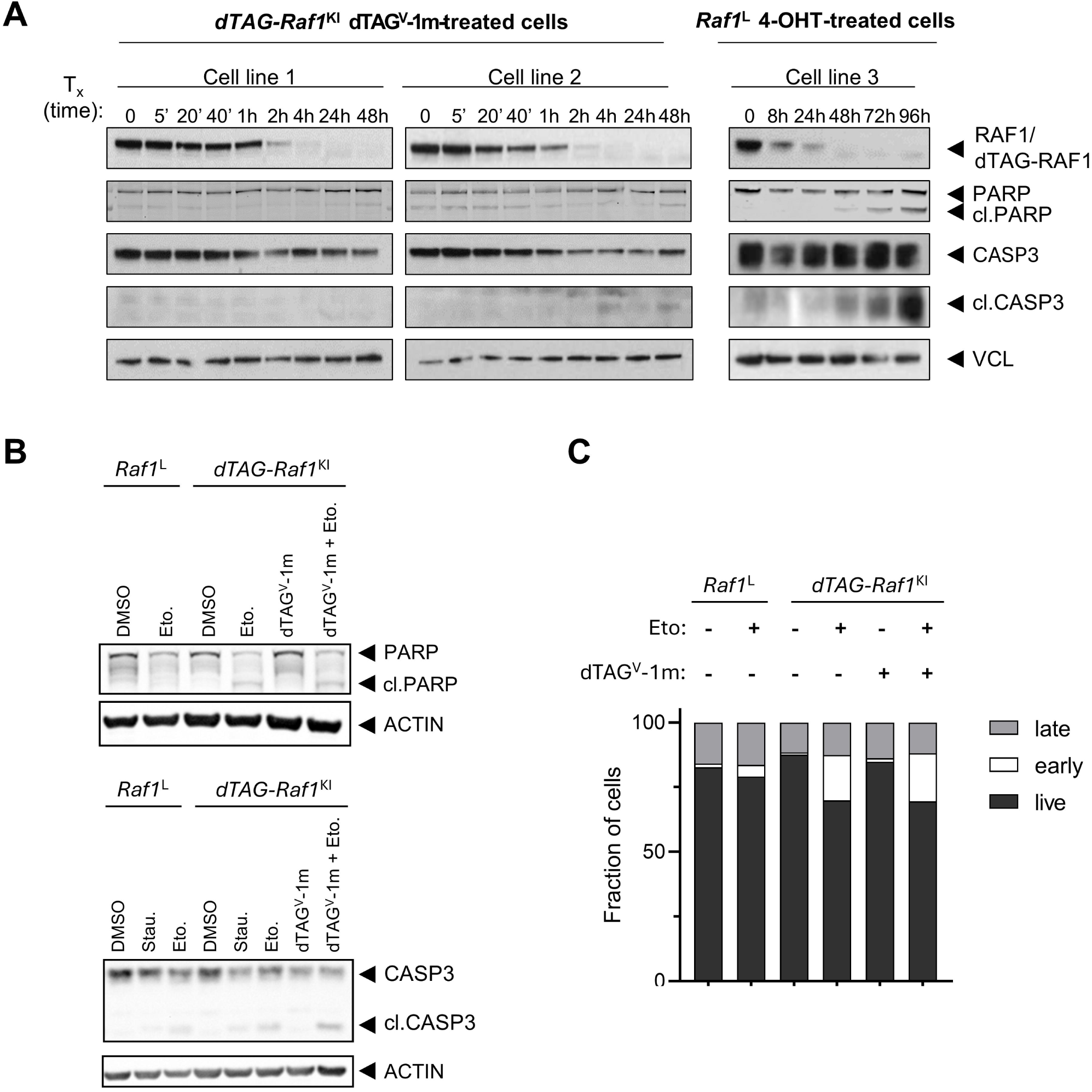
N-terminal dTAG fusion impairs RAF1 scaffold function and apoptotic signaling. (A) Western blot analysis of apoptotic markers following RAF1 pharmacological degradation or genetic *Raf1* elimination. Cells were treated with dTAG^V^-1m or 4-OHT at the indicated time points, according to the genotype. (B) Western blot analysis of apoptotic markers in *Raf1*^L^ and *dTAG-Raf1* cells treated with DMSO, etoposide, dTAG^V^-1m, or the combination of etoposide and dTAG^V^-1m, as indicated. PARP cleavage and caspase-3 cleavage (cl.PARP and cl.CASP3, respectively) were assessed as readouts of apoptotic induction. Actin was used as a loading control. (C) Flow cytometric quantification of apoptosis by Annexin V/PI staining across the indicated conditions. Bars represent the fraction of live (Annexin V□/PI□), early apoptotic (Annexin V□/PI□), and late apoptotic or necrotic (Annexin V□/PI□) cells.

To determine whether the N-terminal dTAG altered the RAF1 interaction landscape, we compared the interactomes of WT RAF1 and dTAG-RAF1 in LUAD-derived cell lines by IP-MS analysis. Although many RAF1-associated proteins were preserved, quantitative comparison identified a subset of interactors with altered abundance in the dTAG-RAF1 pulldown (Fig 5A). Pathway enrichment analysis of these differentially associated proteins revealed a notable overrepresentation of apoptosis and programmed cell death-related pathways (Fig 5B). A more focused pathway analysis using the Reactome database further identified reduced interactions with proteins linked to apoptotic regulation, including ASK1 (Fig 5C), a known RAF1-interacting protein implicated in apoptotic regulation (Chen *et al*, 2001).

**Fig 5.**
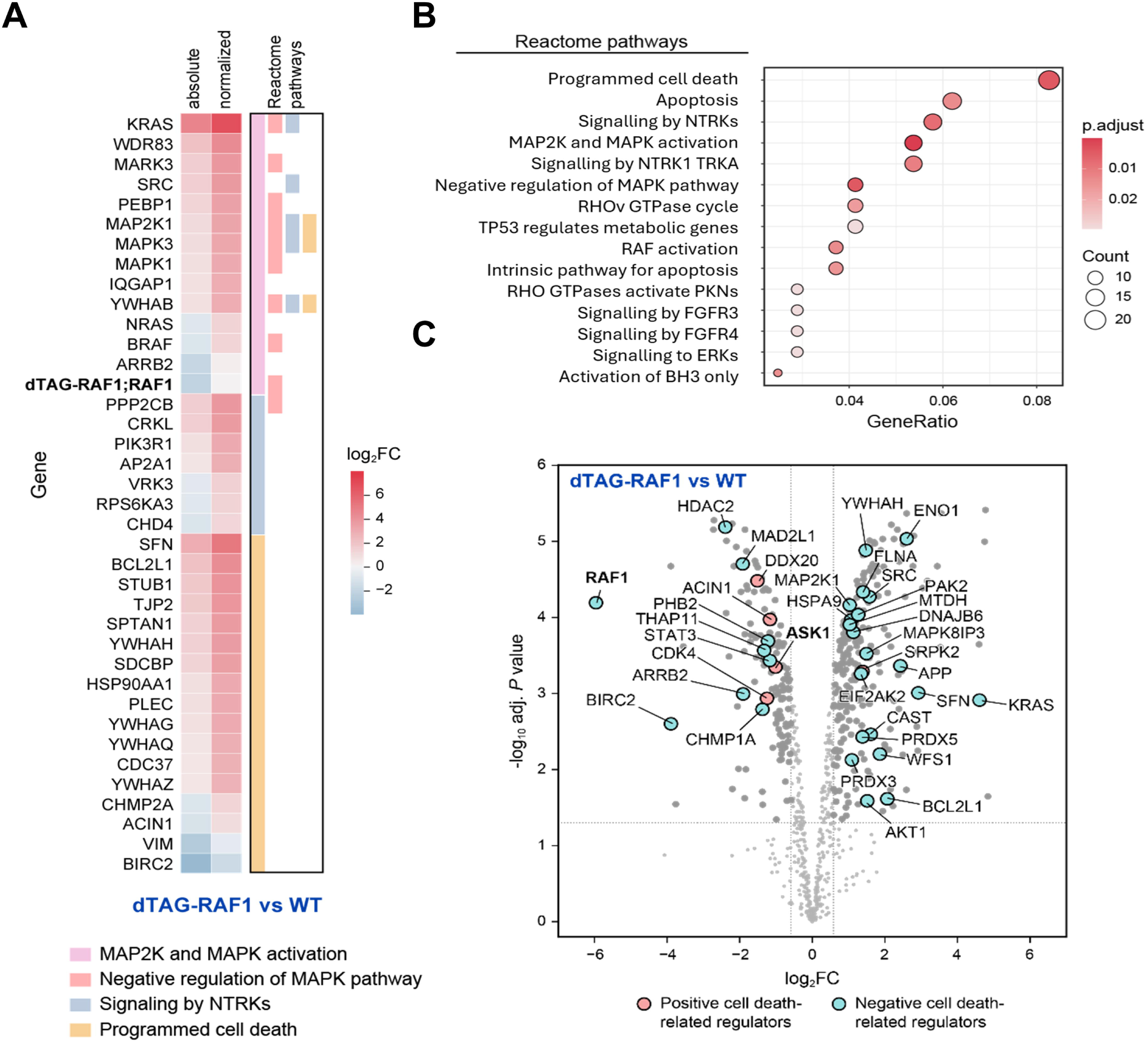
Interactome analysis of dTAG-RAF1 compared to WT RAF1. (A) Heatmap showing RAF1-interacting proteins with increased or decreased association with dTAG-RAF1 relative to WT RAF1. Related cellular pathway is also color-coded as indicated. (B) Top enriched RAF1-interacting pathways observed in dTAG-RAF1 vs WT RAF1-expressing cells (descriptors from the Reactome Pathway database). (C) Volcano plot analysis of positive and negative cell death regulators differentially associated with dTAG-RAF1 relative to WT RAF1.

In contrast, no comparable disruption of MAPK-associated interactions was observed, consistent with the maintained ERK phosphorylation detected *in vivo*. Together, these findings suggest that the N-terminal dTAG does not globally disrupt RAF1 expression or MAPK-associated signaling, but rather alters a subset of RAF1 interactions, including proteins linked to apoptotic regulation.

## Discussion

TPD has emerged as a promising strategy to modulate proteins that remain difficult to inhibit pharmacologically, including multidomain signaling and scaffold proteins. RAF1, a critical effector downstream of KRAS, represents a particularly relevant example in KRAS-driven LUAD, where previous genetic studies demonstrated that *Raf1* ablation promotes robust tumor regression without inducing significant toxicity in healthy tissues (Sanclemente *et al*, 2018). Notably, this tumor-suppressive effect does not result from inhibition of canonical MAPK signaling but instead relies on the disruption of kinase-independent scaffold functions involved in apoptosis regulation. These findings position RAF1 as a useful model to investigate the biological consequences and therapeutic implications of induced protein degradation *in vivo*.

However, targeting RAF1 with small molecules remains challenging. Unlike BRAF, which commonly harbors oncogenic kinase-activating mutations, the RAF1 kinase domain is deeply buried within the HSP90-CDC37 chaperone complex, rendering it sterically inaccessible. Given the absence of selective RAF1 degraders, we employed the dTAG system as a chemogenetic approach to induce acute RAF1 degradation *in vivo*. Our findings demonstrate that efficient depletion of dTAG-RAF1 does not reproduce the tumor regression phenotype previously observed following genetic *Raf1* deletion. Instead, several lines of evidence indicate that the dTAG-RAF1 protein fails to fully replicate the functions of endogenous RAF1. More broadly, these observations highlight an important limitation of chemogenetic degradation systems when applied to scaffold proteins, whose biological activities rely on dynamic and spatially regulated protein-protein interactions that may be particularly sensitive to steric perturbations introduced by fusion tags. Previous studies using chemogenetic degradation systems have reported divergent phenotypes relative to conventional knockout models for proteins including ASH2L, CDK2, and CDK5 (Barsoum *et al*, 2023; Yenerall *et al*, 2023), further highlighting the importance of validating functional equivalence in these degradation platforms.

As a secondary exploratory analysis, we characterized the RAF1 ubiquitination sites within the native RHC complex. Integration of large-scale ubiquitinome datasets (Hansen *et al*, 2021; Rivera *et al*, 2021; Udeshi *et al*, 2020), curated repositories, and structural accessibility analyses identified K462 and K575 as the most solvent-exposed lysine residues within the resolved RHC structure. However, because degrader-induced ubiquitination depends on ternary complex geometry, E3 ligase identity, linker architecture, and induced proximity, these naturally observed or predicted ubiquitination sites should not be interpreted as direct predictors of degrader-productive ubiquitination. Rather, they provide structural context for accessible RAF1 surfaces that may inform future studies.

Systemic dTAG^V^-1m administration in non-tumor-bearing mice was well tolerated, with no significant toxicities observed across physiological, hematological, and histopathological parameters. These findings are consistent with previous genetic ablation studies and further support the notion that RAF1 is dispensable for normal tissue homeostasis in adult mice. Thus, although the dTAG system should not be considered a clinically optimized degrader strategy, these data indicate that systemic RAF1 elimination can be achieved *in vivo* without apparent toxicity.

Despite robust and sustained degradation of dTAG-RAF1 in tumor-bearing mice, this intervention did not reproduce the tumor regression phenotype previously observed following genetic *Raf1* deletion. Tumors continued to grow under dTAG^V^-1m treatment, and acute dTAG-RAF1 degradation did not induce detectable apoptotic markers. These observations indicate that efficient elimination of the tagged protein is not, by itself, sufficient to phenocopy genetic *Raf1* loss in this model.

Several converging findings suggest that this discrepancy reflects functional non-equivalence between dTAG-RAF1 and endogenous RAF1. Even before dTAG^V^-1m treatment, mice expressing the chimeric dTAG-RAF1 protein showed delayed tumor onset and reduced tumor penetrance, suggesting that the N-terminal dTAG partially impairs RAF1-dependent tumor-promoting functions. Although ERK phosphorylation and subcellular localization of dTAG-RAF1 were preserved, quantitative interactome profiling identified a subset of apoptosis-related proteins with altered association in the dTAG context, including ASK1 (Chen *et al*, 2001). Pathway enrichment analysis supported a broader pattern of selective reduction in apoptosis-associated interactors in the dTAG context, while associations with MAPK effectors remained preserved. Together, these findings are consistent with a model in which dTAG tag selectively perturbs RAF1 scaffold-mediated apoptotic regulation without globally disrupting RAF1 expression or canonical MAPK signaling.

Based on these findings, we propose a model in which preserved MAPK-associated signaling and altered apoptotic regulation may account for the inability of dTAG-mediated RAF1 degradation to phenocopy genetic *Raf1* ablation (Fig 6). In the genetic model, *Raf1* deletion induces robust tumor regression associated with increased apoptotic signaling, while canonical MAPK signaling remains unaltered. In contrast, in the dTAG model, MAPK-associated signaling is preserved, but RAF1 scaffold-mediated apoptotic regulation appears to be partially compromised before degrader treatment. As a result, tumors that emerge in the *dTAG-Raf1*^KI/KI^ background may be less dependent on RAF1-mediated apoptotic suppression, limiting the phenotypic consequences of subsequent dTAG-RAF1 degradation. This model illustrates that degrading a functionally compromised protein does not necessarily replicate the effects of genetic ablation.

**Fig 6.**
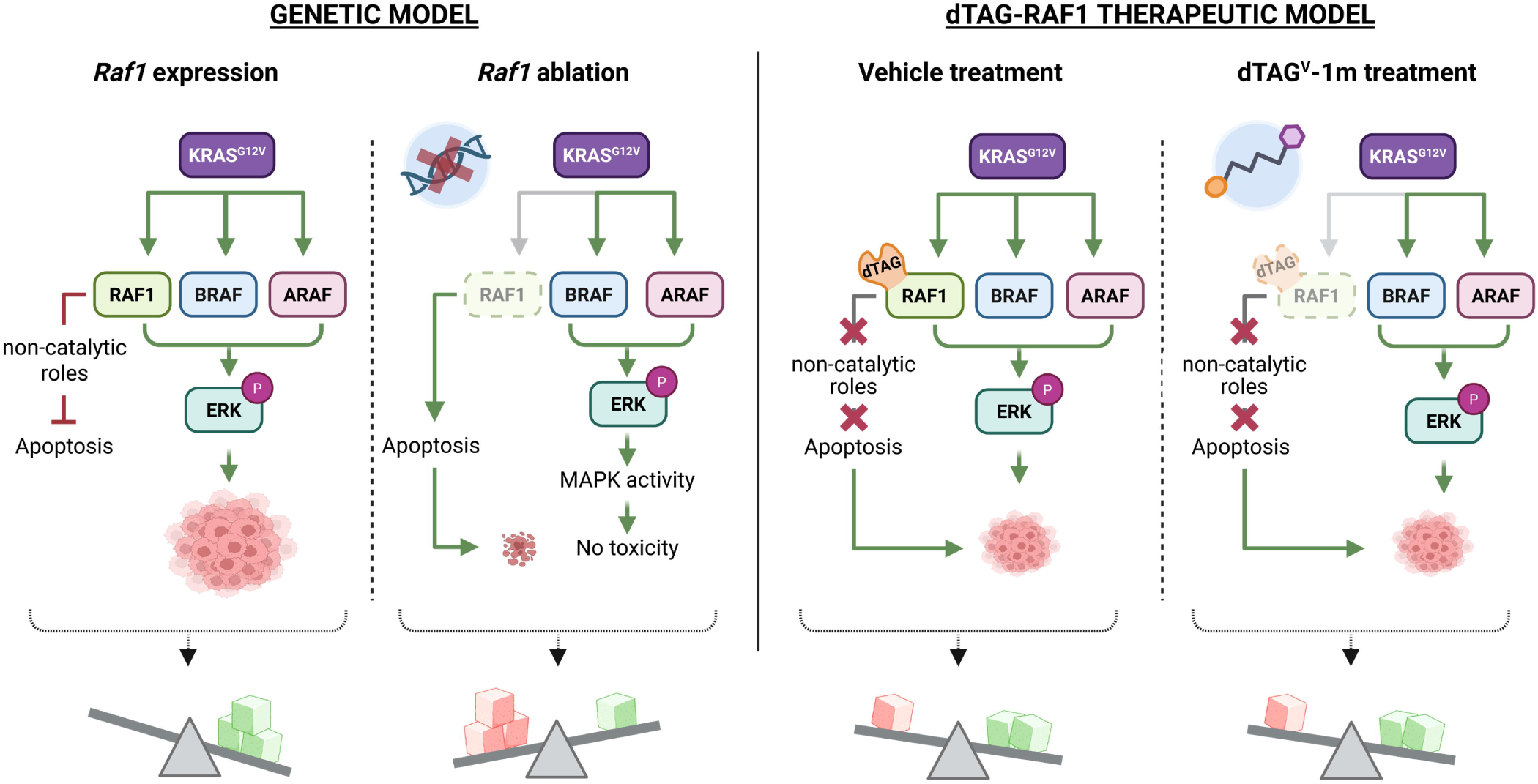
Model for RAF1 function in tumor progression and response to degradation. In *Kras*^G12V^-driven LUAD expressing endogenous RAF1 (*Raf1*^L/L^), canonical MAPK signaling is preserved and RAF1 scaffold functions contribute to suppression of apoptotic signaling, supporting tumor progression. Genetic ablation of RAF1 (*Raf1*^-/-^) disrupts this balance, triggering apoptosis and tumor regression. In tumors expressing the dTAG-RAF1 chimeric protein, MAPK-associated signaling remains intact, but scaffold-mediated apoptotic regulation appears partially impaired, contributing to delayed and reduced tumor progression. Upon dTAG^V^-1m treatment, dTAG-RAF1 is efficiently degraded; however, this does not reproduce the tumor regression observed after genetic *Raf1* ablation. Balance scales illustrate the proposed relationship between proliferative signals (green cubes) and apoptotic regulation (red cubes) in each context.

Beyond RAF1, these findings are relevant to a broader class of scaffolds and regulatory proteins whose biological functions are not fully captured by catalytic activity. For such proteins, degradation may offer advantages over conventional inhibition, but our results underscore that chemogenetic degradation models must preserve the interaction-dependent functions that define target biology. Fusion of chemogenetic tags, while useful for conditional degradation, can disrupt protein-protein interactions and thereby complicate interpretation of target dependency and degradation phenotypes. Accordingly, degrader screening and validation platforms should incorporate structural insights, interactome profiling, and functional readouts, such as apoptosis, differentiation, or tumor regression, in addition to measurements of degradation efficiency or target abundance.

Several limitations of this study should be acknowledged. The tumor treatment cohort was small, mainly due to the reduced tumor penetrance observed in dTAG-Raf1^KI/KI^ mice. Comparisons with historical *Raf1*^L/L^ and *Raf1*^-/-^ cohorts are provided for contextual reference only, as differences in experimental design, treatment regimen, and cohort size preclude direct statistical comparison. Finally, because the dTAG system does not recapitulate the pharmacological properties of a bona fide RAF1 degrader, conclusions regarding the therapeutic potential of RAF1 elimination should be interpreted within this context.

In summary, our study shows that systemic RAF1 elimination using a chemogenetic degradation system is well tolerated *in vivo*, but also reveals that N-terminal FKBP12^F36V^ tag selectively alters RAF1 scaffold-associated functions. Consequently, pharmacological degradation of dTAG-RAF1 does not phenocopy genetic *Raf1* ablation. This work reinforces the concept that, for targets whose tumor-promoting roles are scaffold-mediated rather than catalytic, the fidelity of the degradation tool, not only its efficiency, is a critical determinant of whether chemogenetic models faithfully predict the biological outcome.

## Methods

### Mouse models and ethics

The following genetically engineered alleles were used for this study: *Kras*^LSLG12V^, *Trp53*^L^ (Marino *et al*, 2000), and *dTAG-Raf1*^KI^ (this work). Generation of the *Kras*^LSLG12V^ allele followed procedures similar to those previously described for *Kras*^LSLG12Vgeo^ (Guerra *et al*, 2003).

All mice used in this work were housed in specific-pathogen-free conditions in the Animal Facility of the Spanish National Cancer Research Centre (AAALAC, JRS: dpR 001659), in accordance with Federation of European Laboratory Animal Science Association (FELASA) recommendations and following European Union legislation. Mice were subjected to 12-hours light/dark cycles in ventilated racks under controlled conditions of temperature and humidity. Animals were housed in groups of 4–5 per cage with environmental enrichment. Animals had access to sterilized tap water and chow ad libitum. They were fed with a standardized diet. Both female and male mice were used for the experiments. Health status and body weight were monitored three times per week by the researchers and daily by the animal facility care-takers, ensuring early detection of any signs of distress or disease.

A total of 144 animals were included in the study. 12 tumor-free dTAG-*Raf1*^KI/KI^ mice were used for the systemic tolerability assessment and treated with dTAG^V^-1m (n = 4), vehicle (n = 4), or IGP002 (n = 4) for 30 consecutive days.

For tumor studies, 87 mice were infected intranasally with 1×10^6^ pfu AdCre and 45 with 1×10^7^ pfu. Of these, 8 mice reached the predefined *humane endpoint* before contributing to any analysis and were excluded. Six additional mice from the 1×10^7^ pfu cohort died during or shortly after infection. The remaining mice contributed to tumor onset statistics (Fig 1D-E). Five mice from the 1×10^6^ pfu cohort were used to establish primary LUAD cell lines prior to treatment. Four mice were found dead in cage before treatment initiation, most probably due to respiratory dysfunction. The remaining tumor-bearing mice were randomly assigned to treatment groups: dTAG^V^-1m (n = 7), vehicle (n = 5), or IGP002 (n = 3). IGP002-treated mice were excluded from efficacy analysis following detection of off-target FKBP12 degradation.

Criteria for determining *humane endpoint* included loss of more than 20% of the initial body weight, abnormal activity, abnormal physical appearance (fur, skin, posture, etc.), and signs of respiratory malfunction (dyspnea, rales and extensive atelectasis detected by CT measurement). Once animals showed the above-mentioned symptoms, they were sacrificed within the following 10 minutes. Mice were euthanized by CO_2_ inhalation according to the guidelines of the Ethics and Animal Welfare Committee (CEyBA).

All personnel involved in the study were adequately trained and certified in laboratory animal science, meeting the educational and competency standards recommended by FELASA to ensure responsible and ethical handling of animals throughout the research process. All experiments were approved by the Ethical Committees of the CNIO, the Carlos III Health Institute and the Autonomous Community of Madrid (PROEX 119/17).

### Generation of the dTAG-Raf1 knock-in allele

The endogenous *Raf1* locus was modified using a targeting vector (Gene Bridges GmbH, Heidelberg, Germany) containing the FKBP12^F36V^ coding sequence preceded by a double HA tag, inserted into exon 2. The vector was linearized with AscI and electroporated into G4 ES cells. Recombinant clones were verified by Southern blot using genomic DNA digested with HindIII (left arm) or EcoRV (right arm) and hybridized with external probes located ∼4.5 kb upstream (627 bp) and downstream (589 bp) of exon 2. Three independent embryonic stem (ES) clones were microinjected into C57BL/6J blastocysts to generate chimeric mice. Male chimeras were bred with 129Sv/J females to obtain germline transmission. The NeoR/KanR cassette was excised by fertilizing oocytes from *dTAG-Raf1*^+/KI^ females with sperm from *Tg.CAG-Flp*^T/T^ males. Complete cassette deletion was confirmed by genotyping PCR. All electroporation, microinjection, and genotyping procedures were carried out by the CNIO Transgenic Mice Unit.

### Tumor induction and treatment

Lung adenocarcinomas were induced in 8-to 10-week-old anesthetized mice. Anesthesia was administered via an intraperitoneal injection of ketamine (75 mg/kg) and xylazine (12 mg/kg). Throughout the entire process, the eyes were protected with an ophthalmic ointment (Lacryvisc™) to prevent corneal drying. Tumors were initiated by intranasal instillation of a single dose of 10^6^ plaque-forming units (pfu) of adenovirus expressing optimized CRE recombinase (AdCre). This induced the expression of *Kras*^G12V^ and loss of *Trp53*, leading to multifocal lung adenocarcinomas detectable by CT imaging.

Tumor-bearing mice meeting the inclusion criteria were randomly assigned to treatment groups. CT-based tumor volume quantification was performed blinded to treatment group. Mice with CT-confirmed tumors (≥ 5 mm³) received daily intraperitoneal injections of dTAG^V^-1m (40 mg/kg), synthesized as previously described (Martín *et al*, 2026). The vehicle consisted of 5% DMSO and 20% Kolliphor® EL (Sigma, Cat. 61791-12-6) in saline. The treatment lasted 2 months.

### *In vivo* imaging and tumor volume quantification

Tumor volumes were measured using a SuperArgus CompaCT scanner (Sedecal) under 4% isoflurane anesthesia (Braun Vetcare). Image processing, analysis and 3D rendering were performed using the 3D Slicer Viewer Software. Volumes were calculated assuming ellipsoid geometry:

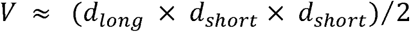

Mice were monitored daily for health status, and CT imaging was repeated every 4 weeks to quantify tumor volume changes. Mice were sacrificed at the end of the experiment or if they reached the *humane endpoint*—loss of more than 20% of the initial weight, anorexia, abnormal activity, abnormal physical appearance, and signs of respiratory distress.

### Systemic tolerability assessment

#### Densitometry

Densitometry scans were performed every 15 days using a Lunar PIXImus densitometer (GE Healthcare) by the Molecular Imaging Unit at CNIO. Mice were anesthetized with 3% isoflurane in oxygen at a flow rate of 0.5 L/min.

#### Echocardiography

Echocardiographic assessments were conducted every 15 days. Anesthesia and thermal regulation were as described for densitometry. A Vevo 3100 micro-ultrasound system (VisualSonics, Toronto, Canada) equipped with a 40 MHz transducer (RMV704) was used. Measurements included the left ventricular internal diameter in systole (LVIDs) and diastole (LVIDd). Left ventricular volumes in systole (LVVs) and diastole (LVVd) were calculated using the Teichholz formula: LVV (mm³) = [7 / (2.4 + LVID)] × LVID³. Ejection Fraction (EF) and Fractional Shortening (FS) were calculated as: EF (%) = [(LVVd − LVVs) / LVVd] × 100; FS (%) = [(LVIDd − LVIDs) / LVIDd] × 100.

#### Hematology

Blood was collected via facial vein (*in vivo*) or cardiac puncture (*ex vivo*). Complete blood counts were performed using a LaserCell blood counter (CVM Diagnóstico Veterinario SL), including total and differential white blood cells, lymphocytes, red blood cells, hemoglobin, and hematocrit.

### Necropsy, histopathology and immunohistochemistry

Necropsies were carried out in the dissection laboratory at CNIO, involving the sacrifice of mice in a CO_2_ chamber (5-min cycle). Death was confirmed by cervical dislocation before tissue harvesting. A comprehensive approach was followed to collect tissue samples, which were then subjected to various preservation methods based on experimental requirements. Specifically, samples were taken in 10% formalin, for subsequent inclusion in paraffin blocks, and cut and stained as needed by the Histopathology Unit at CNIO. Additionally, partial tissue samples were collected in microcentrifuge tubes and snap-frozen in dry ice-cooled 2-methylbutane for the subsequent extraction of DNA, RNA, or protein.

For histopathological visualization, tissue sections of 2.5 μm thickness were stained with Hematoxylin & Eosin (H&E). For immunohistochemistry (IHC) staining, a rat monoclonal α-RAF1 (1:200, Monoclonal Antibodies Unit, Ref. EMI411E) was used. Both H&E and IHC-stained sections were scanned using the Axio Scan.Z1 scanner (Zeiss). Visualization and analysis were performed with QuPath v0.5.1. Histopathological assessment was performed blinded to treatment group.

### Cell culture and RAFless rescue assay

Cell lines used in this study included immortalized *hUBC-CreERT2*^T^*; Araf*^L/Y^; *Braf*^L/L^; *Raf1*^L/L^ (RAFless) MEFs generated in the Barbacid laboratory, and primary LUAD cell lines derived from *Kras*^+/G12V^; *Trp53*^-/-^; *Raf1*^L/L^ (Sanclemente *et al*, 2018) or *Kras*^+/G12V^; *Trp53*^-/-^; dTAG-*Raf1*^KI/KI^ mice established in this work.

Cell lines were maintained at 37°C in a humidified atmosphere containing 5% CO_ and 16% O_ in Dulbecco’s Modified Eagle’s Medium (DMEM; Biowest, Ref. L0103) supplemented with 10% (v/v) fetal bovine serum (FBS; Gibco, Ref. A5256701) and 1% (v/v) Penicillin-Streptomycin (Gibco, Ref. 15140122).

For the RAFless rescue assay, RAFless MEFs were infected with adenoviral particles encoding Cre recombinase to induce genetic ablation of all three endogenous RAF isoforms. RAF ablation was subsequently maintained by supplementing the culture medium with 4-OHT throughout all experiments to prevent re-expression from unrecombined alleles in escaper cells. Then, to assess the ability of the indicated RAF constructs to rescue proliferation in the absence of endogenous RAF proteins, RAFless MEFs were transduced with lentiviral particles encoding WT RAF1, dTAG-RAF1, or empty vector, produced by transient transfection of HEK293T cells using standard packaging plasmids. Following antibiotic selection, 10,000 cells per 100 mm dish were seeded in DMEM supplemented with 10% (v/v) FBS and cultured for 10-14 days with medium replacement every 3-4 days. In cells expressing dTAG-RAF1, dTAG^V^-1m was added at a final concentration of 500 nM and refreshed every two days to maintain sustained protein degradation throughout the assay. At endpoint, colonies were fixed with 4% paraformaldehyde for 15 min at room temperature, washed twice with PBS, and stained with 0.5% (w/v) crystal violet (Sigma).

### Apoptosis induction assays

Cells were treated with either vehicle, 50 µM etoposide, 300 nM dTAG^V^-1m or a combination of both drugs for 24 h. Cells were detached, counted and 200,000 cells per condition were collected in 200 µL of media. Apoptosis was assessed by using the FITC Annexin V Apoptosis Detection Kit I (BD Biosciences #556547), following manufacturer’s instructions. Samples were analyzed by flow cytometry in a Gallios device (Beckman Coulter). Annexin V-FITC and propidium iodide (PI) staining were used to distinguish viable, early apoptotic, and late apoptotic/necrotic cell populations, and data were analyzed and plotted using GraphPad.

For Western blot analysis, cells were treated with either vehicle, 50 µM etoposide, 300 nM dTAG^V^-1m or a combination of both drugs for 24 h. Cell pellets were collected, washed with cold PBS and lysed with 8 M urea containing 1% CHAPS with constant shaking at 4°C for 1 h. Lysates were centrifuged at 13,000 rpm for 10 min at 4°C and the supernatants were collected for protein quantification. For the electrophoresis, 20 μg of protein with loading buffer (NuPAGE LDS Sample Buffer, 10% β-mercaptoethanol) was denatured for 10 min at 70°C, separated on 4-12% SDS-PAGE gels (Invitrogen) and transferred into nitrocellulose membranes using the Trans-Blot Turbo Transfer System. Membranes were blocked in 5% (w/v) non-fat powder milk/TBST or 5% BSA Albumin Fraction V (Panreac Quimica, S.L.U., Ref.A6588.0100) for 1 h and later incubated at 4°C overnight with the primary antibodies in TBS-T. Next day, membranes were washed twice for 5 min in TBS-T and then incubated with the secondary antibodies at RT for 1 h. Enhanced chemiluminescence (ECL) measurement was performed in a ChemiDoc MP Imaging system. The primary antibodies used are the following: β-ACTIN (Sigma, A5441, 1:20,000), PARP (Cell Signaling, 9542, 1:1,000), RAF1 (Cell Signaling, 53745, 1:1,000), CASP3 (Cell Signaling, 9662, 1:1,000). The secondary antibodies used are the following: anti-rabbit (115-035-003) and anti-mouse (111-035-003) from Jackson ImmunoResearch (1:5000 dilution).

### Kinase assay

To assess RAF1-dependent phosphotransferase activity, protein complexes were expressed in Expi293F™ cells by co-transfection of pcDNA3-HA-HSP90 (Addgene, Ref. 22487), pCMV-CDC37-myc-DDK (Origene, Ref. RC201002), and the indicated Strep-tagged RAF1 construct (pcDNA3-2xHA-FKBP12^F36V^-RAF1-Strep, pcDNA3-RAF1-Strep, or pcDNA3-RAF1^Y340E/Y341E^-Strep). 48 hours post-transfection, cells were lysed in buffer containing 20 mM Tris-HCl pH 7.5, 150 mM NaCl, 10 mM MgCl_, 10 mM KCl, 20 mM Na_MoO_, and 0.1% Triton X-100, supplemented with protease and phosphatase inhibitor cocktails and Benzonase. Clarified lysates were purified by affinity chromatography using Strep-Tactin® Sepharose® resin (IBA, Ref. 2-1201-010) and eluted with washing buffer supplemented with 10 mM desthiobiotin (IBA, Ref. 2-1000-002). *In vitro* kinase reactions were performed in ADBI buffer (20 mM MOPS pH 7.2, 5 mM EGTA, 1 mM Na_VO_, 1 mM DTT) supplemented with ATP and Mg²_, using recombinant MEK1 (rMEK; Sigma-Aldrich, Ref. 14-420) as substrate. Constitutively active recombinant RAF1 (rRAF1; Sigma-Aldrich, Ref. R1656) was included as a positive control. Reactions were incubated at 30°C for 30 min and analysed by western blotting.

### Immunofluorescence

Cells were fixed with 4% paraformaldehyde (Thermo Fisher, Ref. 28908) for 10 min at room temperature and permeabilized with 0.2% Triton X-100 in PBS for 10 min. Blocking was performed in PBS containing 1% BSA and 0.1% Tween-20 for 1 hour at room temperature. Primary antibodies α-RAF1 (Santa Cruz, sc-7267, 1:100) and α-HA (Cell Signaling Technology, 3724S, 1:500) were incubated overnight at 4°C, followed by Alexa Fluor 488-conjugated α-mouse (Invitrogen, A11001, 1:250) or Alexa Fluor 647-conjugated α-rabbit (Invitrogen, A31573, 1:250) secondary antibodies for 1 hour at room temperature. Images were acquired using a LIPSI high-content screening system (Nikon) with a 40× objective.

### Western blotting

Protein lysates were prepared in lysis buffer (50 mM Tris-HCl pH 7.5, 150 mM NaCl, 0.5% NP-40) with protease and phosphatase inhibitors. Equal protein amounts (30 µg) were resolved on NuPAGE 4-12% Bis-Tris gels (Invitrogen), transferred to nitrocellulose membranes, and probed with antibodies against: ARAF (Cell Signaling, 4432), BRAF (Santa Cruz, sc-5284), ERK1 (BD Biosciences, 554100), ERK2 (BD Biosciences, 610103), phospho-ERK1/2 (Cell Signaling, 9101), MEK1 (Santa Cruz, sc-6250), MEK2 (BD Biosciences, 610235), phospho-MEK1/2 (Cell Signaling, 9154), RAF1 (BD Biosciences, 610151), Vinculin (Sigma, V9131), CASP3 (Cell Signaling, 9662), cl.CASP3 (Cell Signaling, 9661), and PARP (Cell Signaling, 9542).

### Immunoprecipitation and LC-MS/MS analysis

Cells in 15 cm dishes were lysed in ice-cold IP buffer (50 mM Tris-HCl pH 7.5, 150 mM NaCl, 10% glycerol, 1% Triton X-100, 1 mM EDTA + inhibitors). 200 µg of protein was incubated overnight at 4°C with either α-RAF1 (CST, 53745S) or α-HA magnetic beads (Sigma, SAE0197). Beads were washed and resuspended in 50 mM ammonium bicarbonate for on-bead digestion with trypsin. Peptides were acidified with 1% formic acid, cleaned on C18 stage-tips, fractionated on SCX columns, and reconstituted in 3% ACN, 1% FA for LC-MS.

Peptides were analyzed on an Orbitrap Eclipse Tribrid MS (Thermo Fisher) via an EVOSEP One interface. Separation was performed on a 15 cm × 150 μm C18 column (1.5 μm beads), using an 88-minute DIA gradient (15 samples/day) (Bache *et al*, 2018). MS1 scans (m/z 350-1,200) were acquired at 120,000 resolution; MS2 scans used 33 DIA windows at 30,000 resolution. HCD fragmentation used 28% normalized collision energy. Data were acquired using Tune v3.5.3890 and Xcalibur v4.5.

### Proteomic and interactome analysis

Raw data were analyzed using DIA-NN (v1.8.2 beta 27) in library-free mode (Demichev *et al*, 2020). Searches were run against the *Mus musculus* UniProt database (February 2024), with custom entries for dTAG-RAF1 and common contaminants. Variable modifications included N-terminal acetylation, methionine oxidation, and methionine excision. MaxLFQ-based quantification (Cox *et al*, 2014) was performed via the diann-r package in R (v4.0.1). Proteins were filtered by q-value (< 0.01), peptide count, and missingness. Differential expression was analyzed using limma (Ritchie *et al*, 2015); significance was defined as |FC| > 1.5 and adjusted p < 0.05.

Interactors were filtered based on cytosolic localization (GO and Human Protein Atlas (Thul *et al*, 2017)) and cross-referenced with BioGRID (Oughtred *et al*, 2021) and HuRI (Luck *et al*, 2020) (accessed May 2024). Pathway enrichment was conducted in R using clusterProfiler (Yu *et al*, 2012) (v2023.12.1) with Reactome annotations (Milacic *et al*, 2024) and FDR < 0.05. A secondary enrichment analysis using the Reactome database annotations was restricted to cell death-related terms. Visualizations were generated using enrichplot and heatmaps via seaborn (Waskom, 2021) in Python (v3.12.4).

### Exploratory ubiquitination site compilation and structural annotation

To compile a comprehensive RAF1 ubiquitination profile, data were integrated from three independent sources: experimentally validated sites retrieved from curated repositories (PhosphoSitePlus), computationally predicted sites generated using the RUBI and UbiSite *in silico*, and lysine-GlyGly (K-ε-GG) peptides identified across three large-scale ubiquitinome datasets (Hansen *et al*, 2021; Rivera *et al*, 2021; Udeshi *et al*, 2020).

For structural analysis, the identified lysines were mapped onto available three-dimensional models of RAF1 using PyMOL (v3.0, Schrödinger LLC). Structures used included RAF1 models from the PDB (3OMV and 7Z38). This analysis was used for structural annotation of reported RAF1 lysine ubiquitination sites and was not intended to predict lysines productively modified upon degrader-induced E3 ligase recruitment.

## Statistical analysis

All statistical analyses were performed with GraphPad Prism (v8.4.0) or R (v4.4.2). Data are presented as mean ± SEM. For two-group comparisons, unpaired Student’s t-test (parametric) or Mann-Whitney U test (non-parametric) were used. Longitudinal tumor data were analyzed with linear mixed-effects models fitted by REML, which leverage repeated measurements per animal to improve statistical resolution given the small cohort size, a direct consequence of the reduced tumor penetrance of the *dTAG-Raf1*^KI/KI^ model. Significance was defined as: p < 0.05 (*), < 0.01 (**), < 0.001 (***), < 0.0001 (****).

Sample sizes were estimated based on effect sizes and variances observed in previous studies from this laboratory using the same GEMM (Sanclemente *et al*, 2018, 2021). No formal power calculation was performed.

## Data availability

The mass spectrometry proteomics data generated in this study have been deposited in the ProteomeXchange Consortium via the PRIDE partner repository (Perez-Riverol *et al*, 2022) under accession number PXD079461. All other data supporting the findings of this study are available within the paper and its supplementary information.

## Author contribution

**Laura de-la-Puente-Ovejero:** Conceptualization; Methodology; Investigation; Formal analysis; Data curation; Writing-original draft; Writing-review and editing. **Ana Domostegui:** Investigation; Formal analysis; Data curation; Writing-review and editing. **Inés María García-Pérez:** Investigation; Formal analysis; Writing-review and editing. **Gonzalo Aizpurua:** Investigation; Writing-review and editing. **Lucía Lomba-Riego:** Investigation; Writing-review and editing. **Pilar Ximenez-Embún:** Investigation; Formal analysis; Writing-review and editing. **Marta Isasa:** Investigation; Formal analysis; Writing-review and editing. **Cristina Mayor-Ruiz:** Resources; Supervision; Funding acquisition; Writing-review and editing. **Mariano Barbacid:** Conceptualization; Funding acquisition; Project administration; Writing-review and editing. **Sara García-Alonso:** Conceptualization; Methodology; Formal analysis; Data curation; Supervision; Project administration; Writing-original draft; Writing-review and editing.

## Disclosure and competing interests statement

The C.M-R. lab has received or receives research funding from Aelin Tx and Almirall. C.M-R. is part of the SAB of Cancer State Tx. M.B. and G.A. are co-founders of Vega Oncotargets S.L. G.A. owns 505 shares or 6.58% of the total. M.B. had research contracts with Mirati Therapeutics (2021-2024), Verastem Oncology (2020-2024), Sanofi (2024), and Vega Oncotargets S.L. (2025-2026). M.B. also owns shares of Pfizer (3,241 shares or <0.001%), Bristol Myers Squibb (3,000 shares or <0.001%), Merck Sharp & Dohme (2,184 shares or <0.001%), and Johnson & Johnson (1,042 shares or <0.001%). M.B. was an employee of Bristol Myers Squibb from 1988 to 1999 but he has had no professional contact with this Company since his departure in 1999. The rest of the authors declare no conflict of interest.

## Acknowledgments

The authors would like to express their gratitude to Marta San Román, Alejandra López-García, and Raquel Villar for excellent technical assistance; Isabel Blanco (Animal Facility) and Francisca Mulero (Molecular Imaging Unit) of CNIO for their technical support. This work was supported by the CRIS Cancer Foundation and the Agencia Estatal de Investigación (PID2021-124106OB-I00; MCIU/AEI/10.13039/501100011033). M.B. is a recipient of an Endowed Chair from the AXA Research Fund. M.B. is a recipient of a CIBERONC Fund (CB21/12/00121). S.G-A. was partially supported (from 2019 until 2022) by AECC Postdoctoral Fellowship (call 2020) and by Juan de la Cierva Investigadores fellowship (FJC2018-036013-I). L.d.l.P-O. is supported by a predoctoral fellowship from the Spanish Ministry of Universities (FPU21/04678). I.G-P., G.A. and L.L-R. were supported by a fellowship from “La Caixa” Foundation (LCF/BQ/DR22/11950030, LCF/BQ/DR22/11950011 and LCF/BQ/DR23/12000028). C.M-R. acknowledges funding from the ERC (ERC-2021-StG-101040046 TrickE3), AECC (Cancer Grant Challenges partnership, PROTECT team), and the Spanish Ministry of Science and Innovation (SMSI; PID2023-147995OB-I00, RYC2020-030061-I). A.D. was supported by postdoctoral fellowships FJC2020-043715-I (SMSI) and POSTD246144DOMO (AECC).

## Expanded View Figure Legends

**Fig EV1. N-terminal dTAG fusion partially restores RAF1 function in RAFless cells.** Colony formation assays in RAFless cells transfected with empty vector (pLVX-Empty), dTAG-RAF1 (pLVX-dTAG-*Raf1*), or RAF1-dTAG (pLVX-*Raf1*-dTAG). Cells were treated with DMSO (control) or 4-OHT to eliminate endogenous RAF expression.

**Fig EV2. Characterization of the RAF1 ubiquitination landscape and structural mapping of surface-accessible lysines.** (A) Western blot analysis of RAF1 levels in A549 cells treated with GA alone or in combination with proteasome inhibitors (MG132 or BTZ), both in soluble and insoluble fractions. (B) RAF1 (P04049) peptides identified in SDS-PAGE bands analyzed by mass spectrometry are represented in green. Light and dark shades are used to distinguish different but consecutive peptides in the amino acid sequence. **(C)** Summary of high-confidence ubiquitin sites identified from ubiquitinome repositories, prediction tools, and large-scale datasets. **(D)** Mapping of ubiquitinated lysines onto the cryo-EM structure of the RHC complex (PDB: 7Z38), highlighting the secondary structure context of each resolved lysine. **(E)** Close-up views of the six structurally resolved lysines, colored by solvent-accessible surface area (SASA) score as indicated.

**Fig EV3. Functional characterization of dTAG-RAF1.** (A) Representative images of colony formation in cells treated with DMSO or dTAG^V^-1m for 15 days. (B) Western blot analysis confirming chimeric protein degradation in the same cells at endpoint. (C) Colony formation assays in RAFless cells transfected with empty vector, RAF1 or the chimeric dTAG-RAF1. Cells were exposed to DMSO or 4-OHT to eliminate endogenous RAFs expression. In the case of dTAG-RAF1 treated cells, dTAG^V^-1m was added at 500 nM and refreshed every two days. (D) *In vitro* kinase assay using recombinant MEK1 (rMEK) as a substrate. rRAF1 refers to a commercially available, constitutively active recombinant RAF1 (MW = 60 kDa; not detected in the RAF1 standard blot due to its lower molecular weight). R** denotes the RAF1^Y340E/Y341E^ constitutively active mutant. dRHC refers to the dTAG–RAF1–HSP90–CDC37 complex. (E) Confocal microscopy images illustrating subcellular localization. Scale bar: 20 μm. dTAG^V^-1m: 0.3 μM for 24 hours.

## Notes

### Summary of Updates

The author list has been corrected

