## Supplementary figures and images for "RAF1 scaffold integrity shapes chemogenetic degradation outcomes in KRAS-driven lung cancer"

### Figure EV1

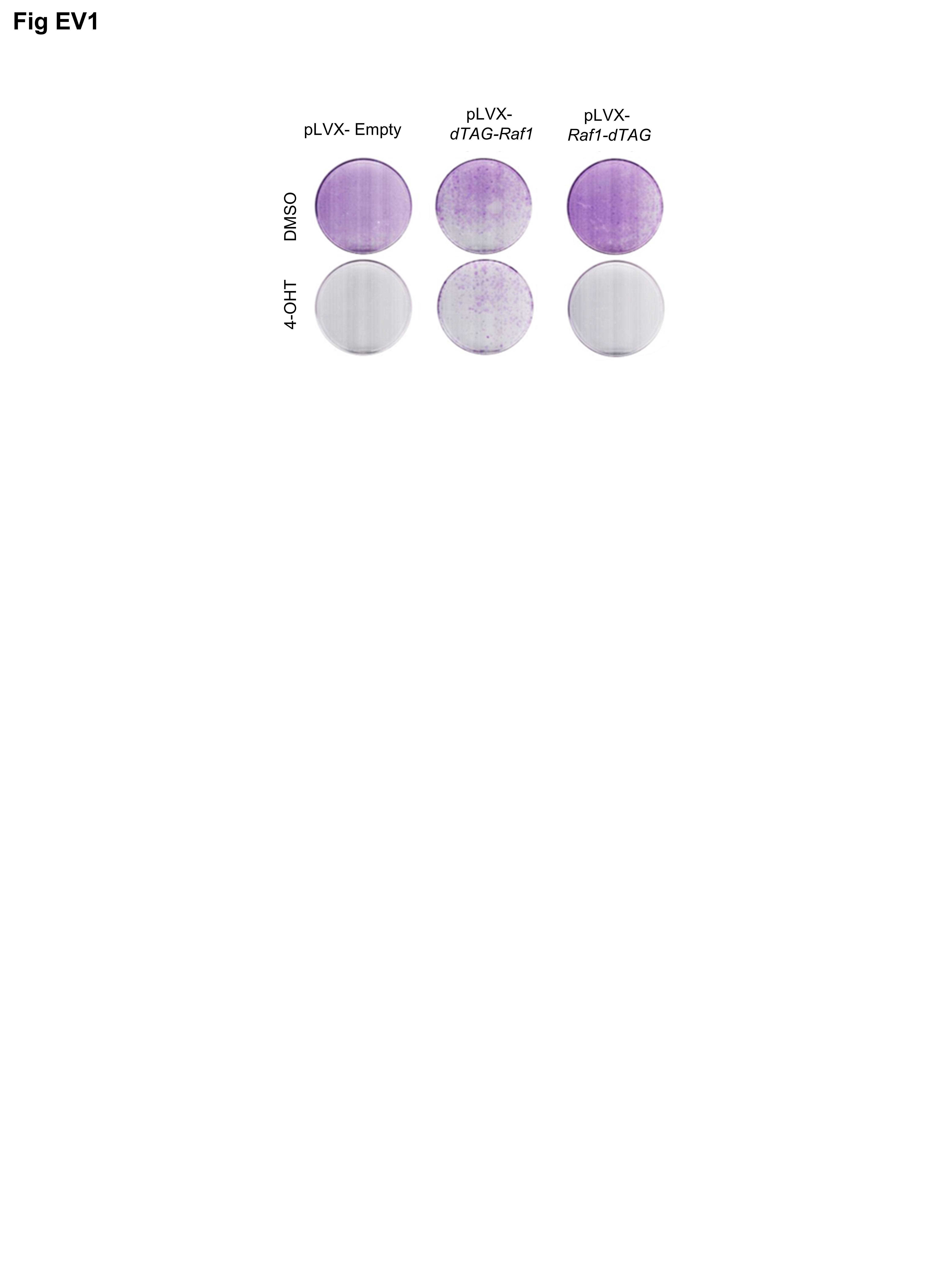

### Figure EV2

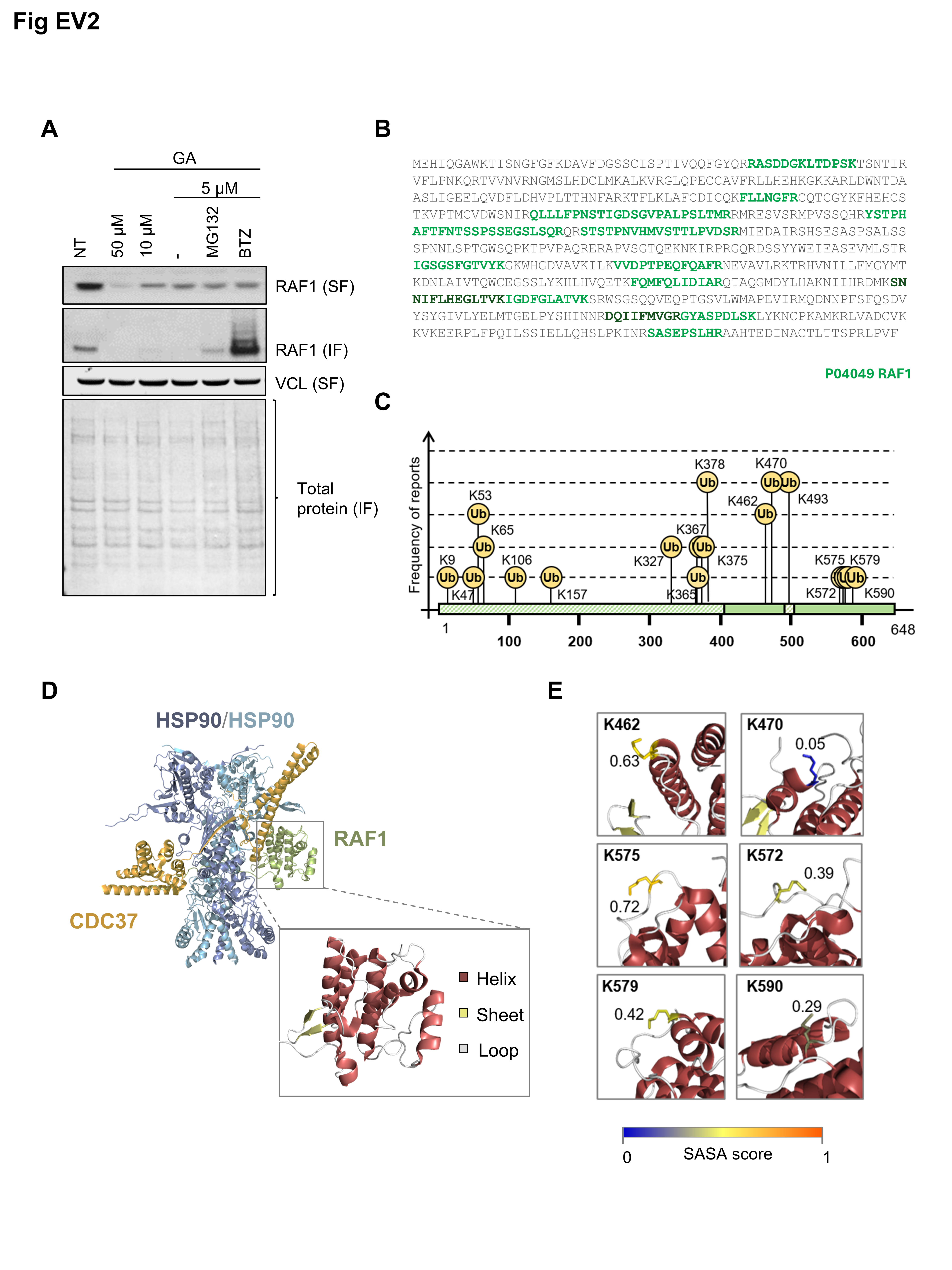

### Figure EV3

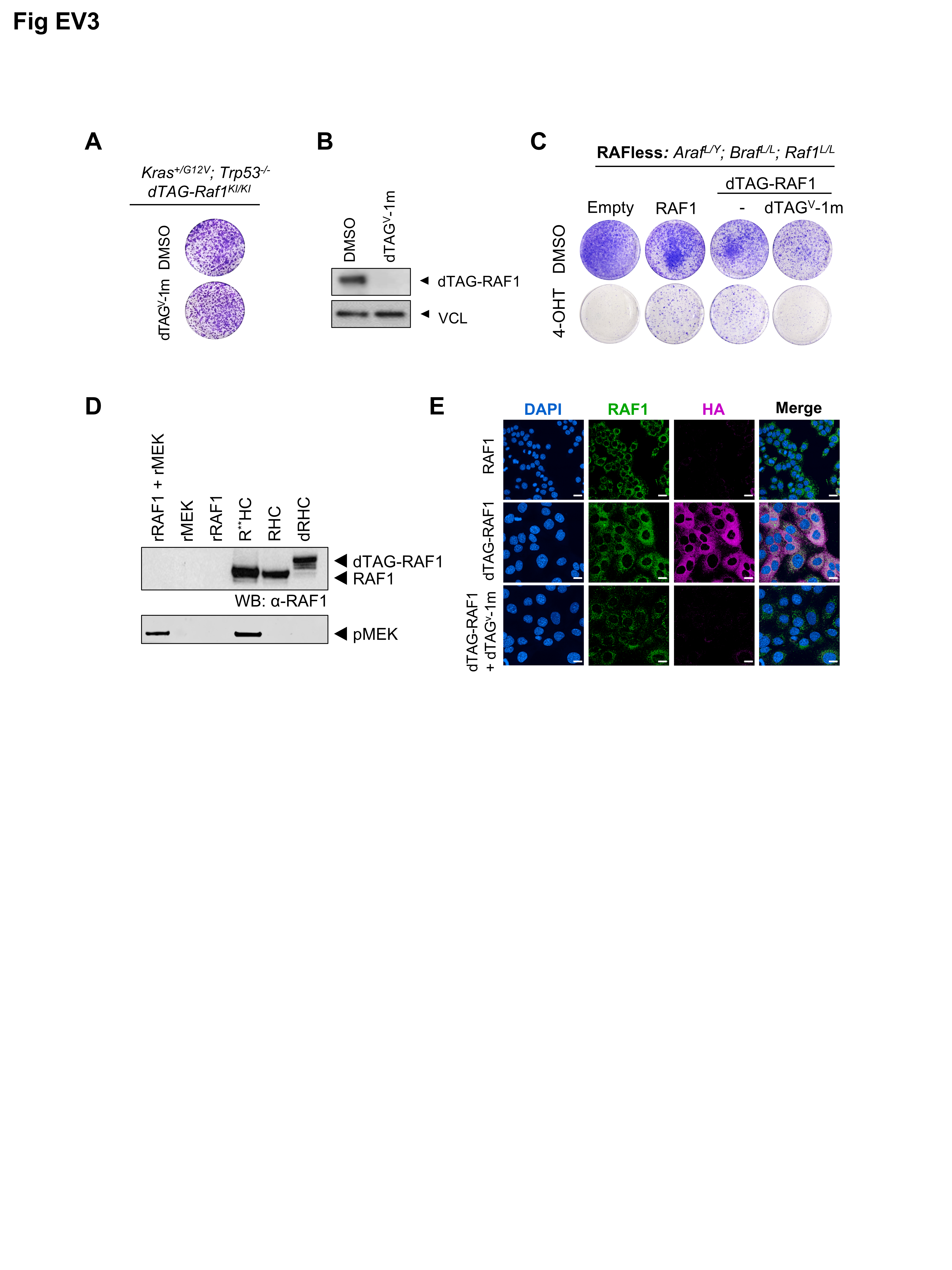
